# Dissecting the energetic architecture within an RNA tertiary structural motif via high-throughput thermodynamic measurements

**DOI:** 10.1101/2022.12.15.518780

**Authors:** John H Shin, Steve L Bonilla, Sarah K Denny, William J Greenleaf, Daniel Herschlag

## Abstract

Structured RNAs and RNA/protein complexes perform critical cellular functions. They often contain structurally conserved tertiary contact “motifs,” whose occurrence simplifies the RNA folding landscape. Prior studies have focused on the conformational and energetic modularity of intact motifs. Here we turn to the dissection of one common motif, the 11nt receptor (11ntR), using quantitative analysis of RNA on a massively parallel array (RNA-MaP) to measure the binding of all single and double 11ntR mutants to GAAA and GUAA tetraloops, thereby probing the energetic architecture of the motif. While the 11ntR behaves as a motif, its cooperativity is not absolute. Instead, we uncovered a gradient from high cooperativity amongst base paired and neighboring residues to additivity between distant residues. As expected, substitutions at residues in direct contact with the GAAA tetraloop resulted in the largest decreases to binding, and energetic penalties of mutations were substantially smaller for binding to the alternate GUAA tetraloop, which lacks tertiary contacts present with the canonical GAAA tetraloop. However, we found that the energetic consequences of base partner substitutions are not, in general, simply described by base pair type or isostericity. We also found exceptions to the previously established stability-abundance relationship for 11ntR sequence variants. These findings of “exceptions to the rule” highlight the power of systematic high-throughput approaches to uncover novel variants for future study in addition to providing an energetic map of a functional RNA.

**Significance Statement:** Properly folded RNAs perform essential biological processes. Many RNAs contain tertiary contact “motifs” whose structural and energetic properties are conserved across different RNAs. This study delves into the nature of an RNA motif. We determined the effects of mutations on the energetic properties of a common tertiary contact motif, referred to as the 11-nucleotide receptor. As deep study of the energetic architecture of this and other RNAs requires thermodynamic measurements for many sequence variants, we used RNA-MaP, a quantitative, high-throughput method, to obtain binding free energies for all single and double mutants of the motif. Our results revealed the energetic architecture of this motif and identified rare variants with unexpected properties.

## Introduction

Folded RNAs provide crucial functions to sustain life, including the maturation of mRNAs, translation of proteins, and regulation of gene expression (1–4). Similar to proteins, these RNAs must traverse a sequence-encoded conformational landscape to fold into functional structures. However, these landscapes differ considerably. For proteins, secondary structure is typically not stable without tertiary structure, and folded globular proteins contain extensive networks of tertiary contacts (5–7). In contrast, stable secondary structure elements allow RNA to fold hierarchically, where preformed secondary structures are brought together by tertiary contacts to form a fully folded, functional RNA (3, 8–10). Also, unlike the case with proteins, tertiary contacts are typically sparse within a folded RNA (Fig. 1*A*; (4)). Comparisons of RNA tertiary interactions have revealed, in many cases, common sequence and structural elements across different RNAs; these elements have been referred to as “RNA motifs” (11–19). These motifs also exhibit conserved energetic behavior in different structural contexts, as described by an RNA Reconstitution Model that builds up the energetics of RNA molecules from the energetics and conformational preferences of their constituent motifs (10, 20, 21). In contrast, we know little about the energetic properties within the motifs themselves — information that defines their cooperativity and evolvability. Here, we explore the energetics of one common tertiary contact motif, the 11-nucleotide receptor (11ntR).

**Fig. 1.**
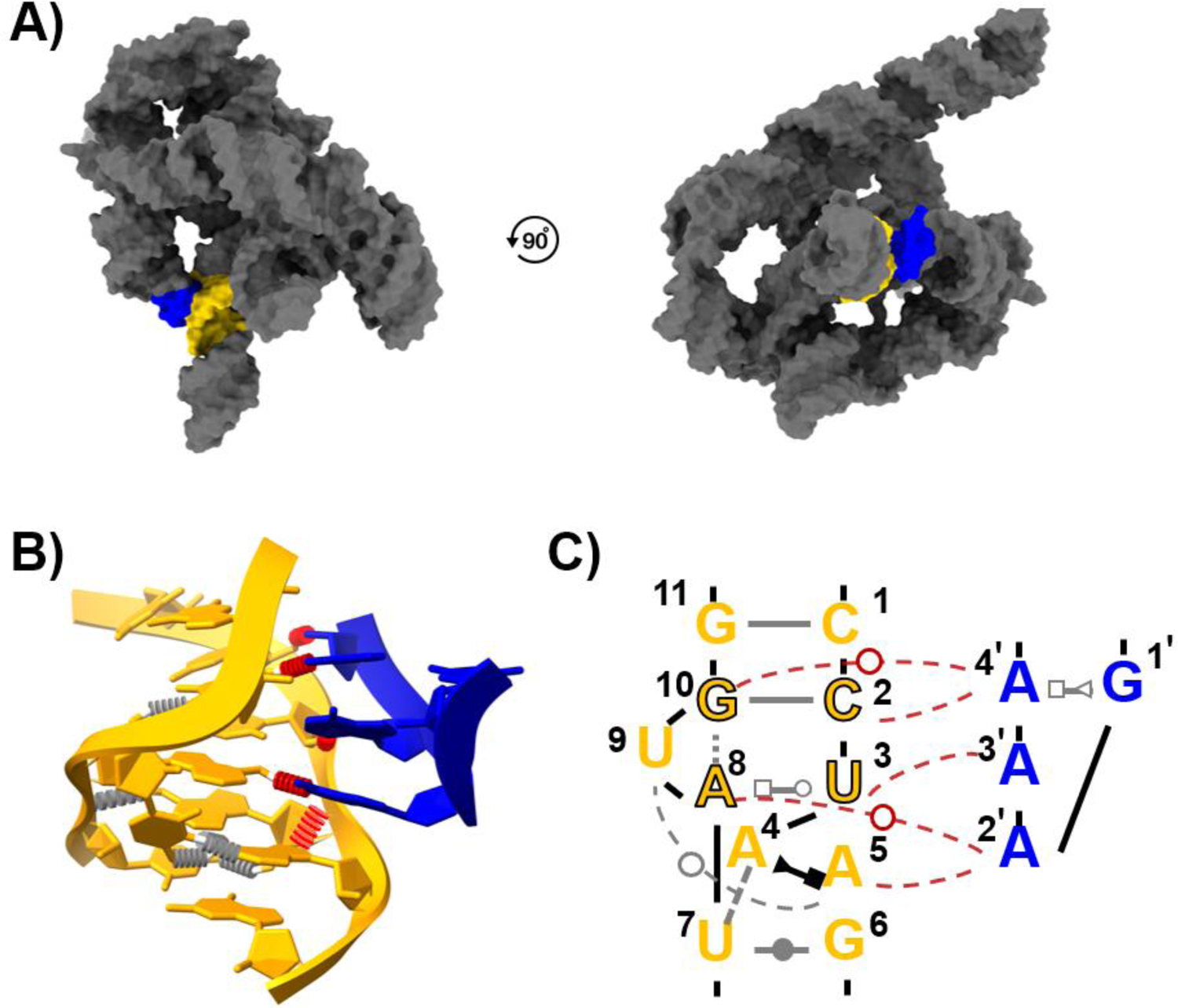
RNA folds via conserved tertiary motifs. (A) Structured RNAs consist of helical segments with interspaced junctions that are joined by sparse tertiary contacts, for example the 11ntR within the Tetrahymena group 1 Intron (PDB ID: 1GID (23), visualized via ChimeraX (66)). (B) The crystal structure of the 11ntR • GAAA interaction motif (11ntR in yellow and GAAA tetraloop in blue). Tertiary interactions are shown in red, and interactions within the 11ntR are shown in gray (base pairing interactions omitted for clarity). (C) Schematic of the 11ntR • GAAA interaction, color coded as in (B). The residue numbering scheme for the 11ntR is taken from Bonilla *et al*. (25) and is used throughout; we number the tetraloop residues 1ʹ to 4ʹ to distinguish them from the receptor residues. Base pairing interactions are denoted using the standard Leontis-Westhof notation in gray (67), and the “core” residues making specific hydrogen-bonding interactions with the tetraloop are outlined in black. Solid lines indicate base-pairing interactions, and dashed lines indicate interactions outside a residue’s base pairing partner.

The 11ntR is highly specific for binding to the GAAA tetraloop and is found in many RNAs (Fig. 1*B* and *C*; (13, 14, 17, 21–32)). Extensive prior studies have found that mutating receptor residues that directly contact the tetraloop result in large decreases in the free energy of binding; mutations to motif residues not directly contacting the tetraloop also reduce binding, but to lesser extents (33). The development of high-throughput methods that yield quantitative thermodynamic measurements over a wide dynamic range provides the opportunity to probe the energetics of the interactions and interconnections within the 11ntR.

Greenleaf and coworkers recently described RNA on a Massively Parallel array (RNA-MaP), which measures thermodynamic and kinetic parameters for tens of thousands of RNA sequences of interest (34, 35). RNA-MaP has been applied to several RNA and RNA • protein systems (36– 41). Pertinent to this study, the high-throughput nature of RNA-MaP enabled exploration of thousands of tetraloop receptors (TLRs) in different local structural contexts, as described in Bonilla *et al*. (40). In brief, Bonilla *et al*. embedded a library of 1493 TLRs into different tectoRNA scaffolds that differ in length and sequence between the two ends of the tectoRNA construct and measured binding free energies (Δ*G*_bind_) (Fig. 2*A* and *B*; (33, 42, 43)). By comparing Δ*G*_bind_ values of tectoRNA constructs containing the same TLRs in different scaffolds, “thermodynamic fingerprints” could be obtained for the TLRs, providing information about the TLR • tetraloop binding conformational preference. Specifically, the difference in Δ*G*_bind_, ΔΔ*G*, is expected to be the same across scaffolds for two TLR variants that share conformational preferences and deviate when the overall motif conformations differ (Fig. 2*C*). Bonilla *et al*. found that the stability of natural 11ntR variants correlated with their frequency in natural RNAs and that they had the same fingerprints as the wild-type, suggesting that they evolved to bind strongly to the GAAA tetraloop with a conserved conformation.

**Fig. 2.**
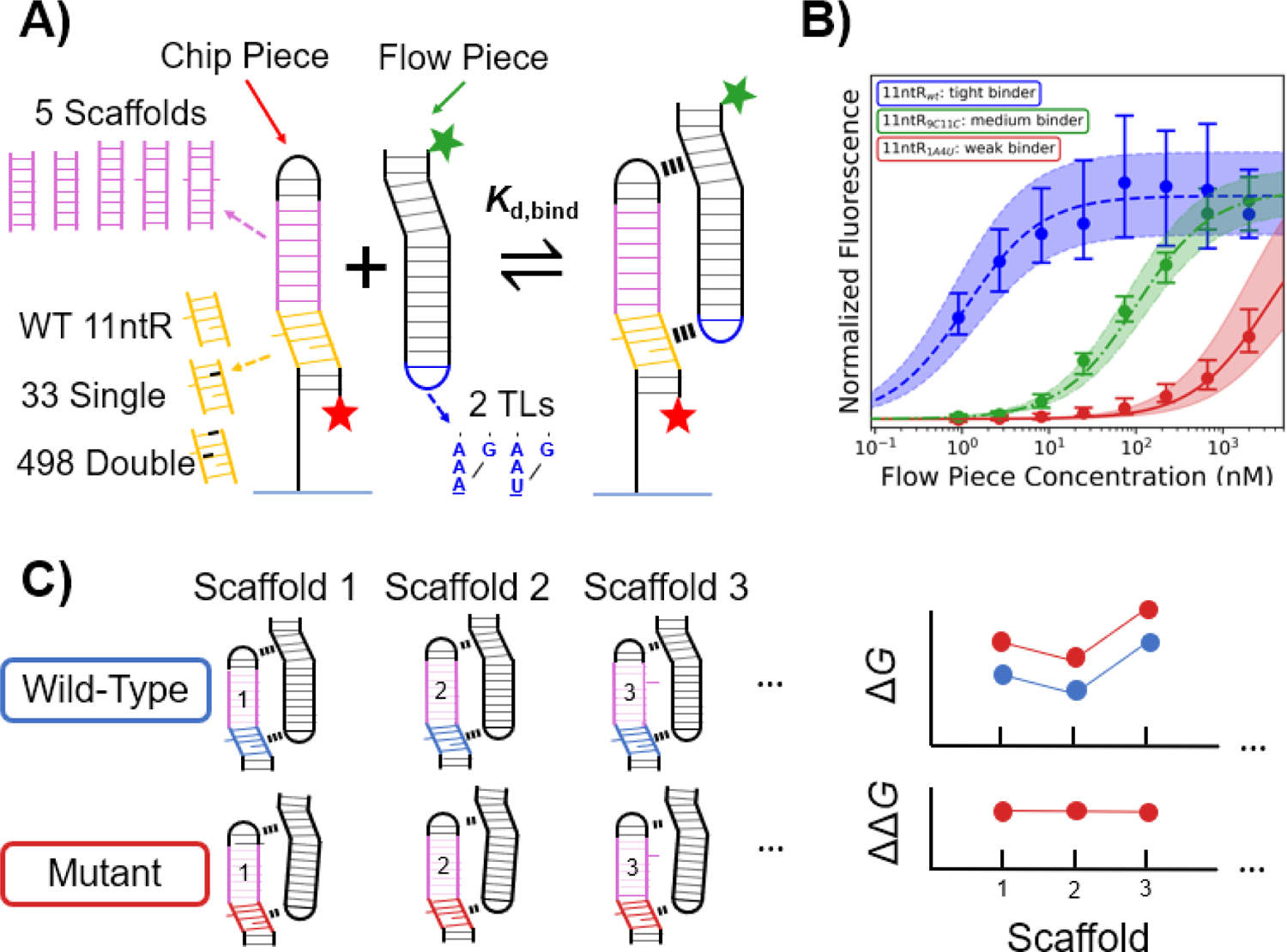
Thermodynamic characterization of the wild-type 11ntR and the 33 single and 495 double mutants of 11ntR via RNA-MaP. (A) Binding for 11ntR mutants to a GAAA and GUAA tetraloop (blue) were measured for five tectoRNA “scaffolds” (pink), consisting of an immobilized chip piece (left) and a free-flowing flow piece (right) (36, 40, 42, 43). (B) Fluorescence measurements across a series of flow piece concentrations were fit to equilibrium isotherms. Fluorescence *versus* concentration (log scale) are shown for three sequence variants: the wild-type 11ntR (Δ*G*_bind_ = – 12.1 kcal/mol (–12.3, –11.9), n = 42 clusters), the double mutant 9C11C (Δ*G*_bind_ = –9.4 kcal/mol, (–9.6, –9.3), n = 70 clusters), and the double mutant 1A4U (Δ*G*_bind_ = –7.4 kcal/mol, (–7.7, –7.1), n = 97 clusters). Data points represent median fluorescence values with 95% bounds, and the isotherms are denoted by the best-fit line running through them. (C) Schematic depicting thermodynamic fingerprints via comparisons using different scaffolds. In this example, the fingerprints of two 11ntR variants, a the wild-type (blue) and a mutant (red), are compared using thermodynamic fingerprints derived with three scaffolds. The Δ*G* values across scaffolds (top plots) are the thermodynamic fingerprints for the two 11ntRs; visual inspection suggests that the fingerprints are identical, and the ΔΔ*G* values are the same across the scaffolds for this example, *i*.*e*., there is zero variance in the ΔΔ*G* values. Thus, we conclude that the mutant shares the same fingerprint as the wild-type. In actuality, we determine if the variance is greater than predicted for experimental error.

In this work, we deeply analyze a sub-library from Bonilla *et al*. (40) designed to dissect the energetic architecture of the 11ntR. This sub-library encompasses the wild-type 11ntR as well as all single and double mutants thereof binding to two tetraloops, the cognate GAAA loop and an alternate GUAA loop, which binds much weaker to the 11ntR (Fig. 2*A*; (13, 26, 33, 40)). The results reveal a gradient of energetic effects from mutations throughout the motif, show that the energetic interconnectivity is reduced with the weaker-binding GUAA tetraloop, and uncover unexpected patterns of energetic cooperativity, rescue, and synergy within the motif. Our deep investigation into the sequence-energy landscape of the 11ntR also identified a small number of sequence variants that form stable 11ntR • GAAA interactions but are not observed in nature. Extensive and thermodynamically accurate data from RNA-MaP and other emerging platforms will extend and deepen our understanding of the relationships between sequence, structure, and energetics, how nature has utilized these conformational and energetic landscapes for evolution of RNA motifs, and how we can exploit this information to engineer and precisely tune novel functional RNAs.

## Results and Discussion

### Curation of the 11ntR thermodynamic data

We analyzed 5290 equilibrium dimerization measurements of tectoRNAs containing 11ntR variants from a parent RNA-MaP dataset from Bonilla *et al*. (40). Our 11ntR library consisted of a wild-type sequence along with all 33 single mutants and all 495 double mutants, for a total of 529 variants (Fig. 2*A*). Each variant was placed in the context of five different tectoRNA “scaffolds” and binding was measured to a partner tectoRNA monomer with a GAAA or GUAA tetraloop (Fig. 2*A* and *B*, *SI Appendix* Fig. S1). We only report *K*_d_ (and Δ*G*_bind_ = –*RT* ln *K*_d_) values for variant-scaffold combinations with more than five molecular repeats on the RNA MaP array (clusters) to ensure high-quality data (36, 40); 95% of the sequences had more than five clusters in the GAAA dataset with a mean of 65 clusters per sequences, and 91% of sequences had more than five clusters in the GUAA dataset with a mean of 32 (*SI Appendix* Fig. S2*A* and *B*). We obtained at least one binding measurement for each 11ntR variant across the scaffolds; 423 and 417 of the 529 variants gave data for all five of the scaffolds when binding to the GAAA and GUAA tetraloops, respectively (*SI Appendix* Fig. S2*C* and *D*). We discuss the data in terms of ΔΔ*G*, the difference in Δ*G*_bind_ between 11ntR variants and the wild-type 11ntR, such that greater destabilization leads to more positive ΔΔ*G* values. Background fluorescence of the tectoRNA at high concentrations precludes measurement of Δ*G*_bind_ values weaker (less negative) than −7.1 kcal/mol, so variants with a Δ*G*_bind_ value above this threshold are reported as limits rather than explicit values. We refit the fluorescence data presented in Bonilla *et al*. (40) using larger bootstrap resampling (*n* = 10,000) of Δ*G*_bind_ values to strengthen statistical tests, as described in the *Methods*. The refitting procedure did not significantly change the Δ*G*_bind_ values measured for the variants presented herein (*SI Appendix* Fig. S3).

Our dataset of 529 11ntR variants contains 12 variants previously characterized by Geary *et al*. (33), and we find good agreement with their prior thermodynamic results (*ρ* = 0.96; *SI Appendix* Fig. S4). The measurements were performed using different tectoRNA scaffolds and ionic conditions, suggesting that the energetic effects of the 11ntR motif are independent of these external factors. This additivity in energetics gives additional support to the Reconstitution Model of RNA folding (10, 20).

### Single mutations to the 11ntR give a gradient of deleterious effects

Structures of RNA molecules containing the 11ntR • GAAA interaction reveal a core set of residues that make direct contacts between the tetraloop and the receptor (Fig. 1*C*; (23, 28)). These interactions are represented schematically as red dashed lines in Fig. 3*A*: an A-minor motif between the G10-C2 pair (blue) and the tetraloop residue A4ʹ; a “UA-handle” consisting of hydrogen bonds between the A8-U3 pair (orange) and tetraloop residues A2ʹ and A3ʹ; and stacking between A4-A5 dinucleotide platform (red) and A2ʹ in the tetraloop. A naive expectation would be that mutations to groups making direct interactions have more deleterious effects on binding (larger ΔΔ*G*) than mutations outside the core where these interactions are made. Deleterious effects from mutations to residues outside of this core would presumably arise from local rearrangements that propagate through the structure to sites of direct interaction.

**Fig. 3.**
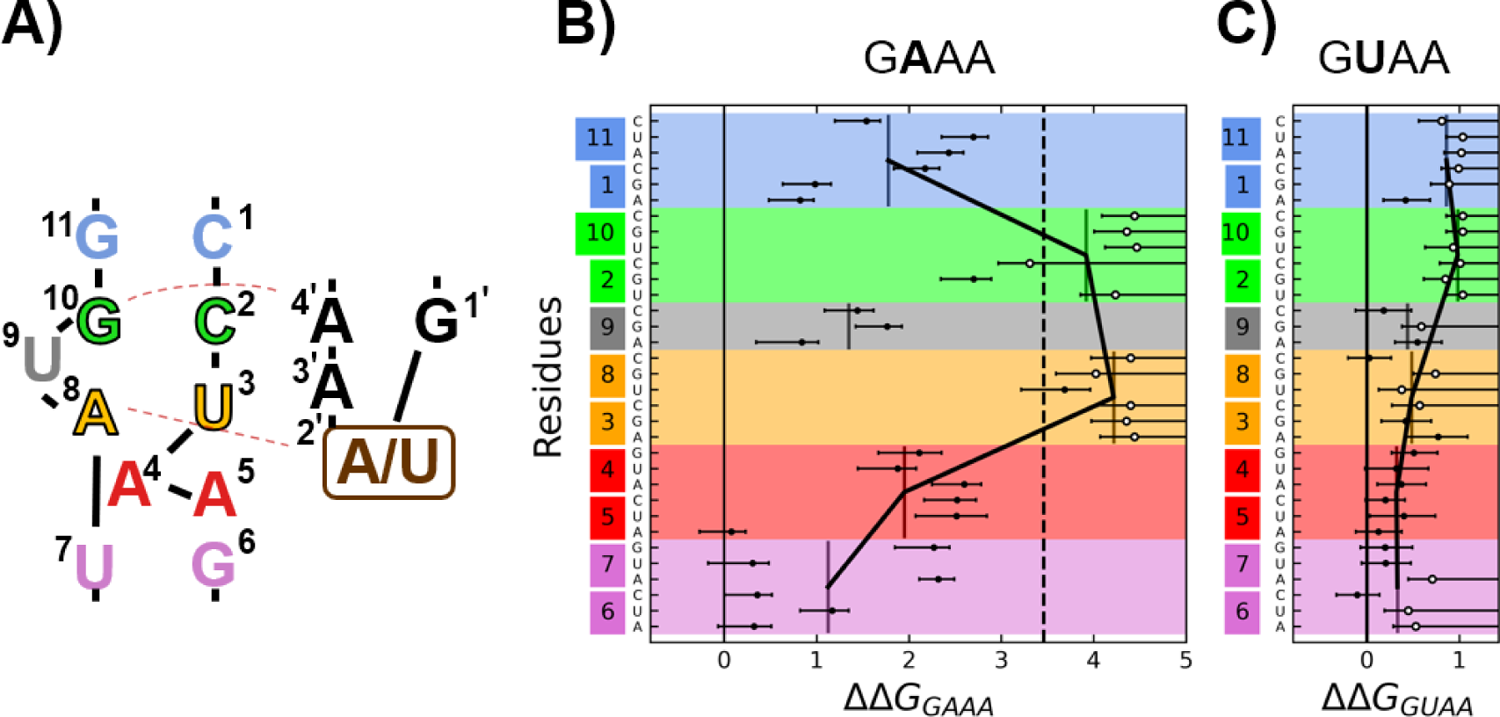
Single mutations to the 11ntR have larger effects in the core and with the cognate GAAA tetraloop receptor. (A) The 11ntR consists of five sets of stacked base pairs (blue, green, yellow, red, and purple) and an additional bulged base at position 9 (gray). Hydrogen bonding interactions (excluding base-phosphate interactions) between the 11ntR and GAAA are shown as dashed red lines, and these interactions occur with the “core” residues in the 11ntR outlined in black. The ΔΔ*G* values for binding to GAAA (B) and GUAA (C) of single mutants (points) relative to the wild-type 11ntR. The average ΔΔ*G* per residue (vertical lines) are plotted grouped by base pair (y-axis colors), and the average points for the five base steps are connected by black lines. In (B), the ΔΔ*G* of the wild-type 11ntR binding to GUAA versus GAAA is shown as the dashed line. Open circles represent lower limits of ΔΔ*G* values, and error bars represent 95% confidence intervals for the ΔΔ*G* values (limits have no upper bounds on confidence intervals).

The single-mutation results, summarized in Fig. 3*B*, support both expectations. On average, mutations to the core (green/orange) have the largest effects, residues stacked against the core (blue, red) have smaller effects, and the remaining residues have the smallest effects (gray, purple). No mutation to a residue outside the core results in a larger ΔΔ*G* than those within, consistent with peripheral residues helping to position the core residues to interact with the GAAA tetraloop rather than making direct interactions. U9 (gray) is flipped out of the helical stack and hydrogen bonds to the A4-A5 platform but not the tetraloop (Fig. 1*C*). Thus, we expect its mutational effects to be more modest than for the core residues that surround it, and that is indeed the case (ΔΔ*G*_9_ < 2 kcal/mol *versus* ΔΔ*G*_core_ > 2.5 kcal/mol). Mutations at base step U7-G6, which makes no direct interactions to the tetraloop but stacks against the A4-A5 dinucleotide platform, gave either negligible effects or an effect that is similar to that of deleterious A4-A5 mutations (Fig. 3*B*, purple and red). Presumably, the U7-G6 step positions A4-A5 to stack against the tetraloop; the large fraction of U7-G6 mutations that give no or negligible effects suggests that there is flexibility in the ability to form the A4-A5 • A2ʹ stacking interaction. Additionally, we found no evidence of a change in binding conformation from single mutations (*i*.*e*., no significantly negative correlations of the Δ*G* values across scaffolds between single mutants and the wild-type 11ntR binding to GAAA; *SI Appendix* Fig. S5*A*), suggesting that these changes in binding energy come from local conformational changes that do not impact the overall orientation of the mutant 11ntR • GAAA interactions.

We next analyzed binding data for the wild-type 11ntR with a G**U**AA tetraloop instead of the canonical G**A**AA tetraloop. Our results show much weaker binding of the 11ntR to the G**U**AA tetraloop (relative to G**A**AA), in agreement with prior data (ΔΔ*G* = 3.5 kcal/mol, black dashed line in Fig. 3*B*; (13, 40)). This large effect presumably arises because A2ʹ of the tetraloop, which makes multiple direct interactions with the 11ntR, is mutated to a U, thereby removing several direct tertiary interactions (Fig. 1*B* and *C*). Nevertheless, the thermodynamic fingerprints across the five scaffolds are not significantly different between the two tetraloops (*SI Appendix* Fig. S5*B*-*D*), suggesting that the overall geometry of the tertiary contact is the same despite the loss of interactions and large decrease in binding energy.

### Removal of the interacting A_2_ residue of the GAAA tetraloop results in greatly reduced energetic penalties from core 11ntR mutations

Given the expected loss of interactions between the wild-type 11ntR and the G**U**AA tetraloop *versus* the G**A**AA tetraloop, we predicted that 11ntR mutations would be less deleterious for G**U**AA than for G**A**AA tetraloop binding. Specifically, mutating residues that position the 11ntR to interact with A2ʹ of the G**A**AA tetraloop is predicted to have smaller effects to binding. While the weaker overall binding precludes measurements for many of the mutants, the measurable Δ*G*_bind_ supported this expectation: the binding free energy effects in Fig. 3*C* (G**U**AA) are smaller than those in Fig. 3*B* (G**A**AA) as seen by the magnitude of ΔΔ*G* values and the shallower dependence of ΔΔ*G* values on the base step (line plots in Fig. 3*B* and *C*).

We first looked at the residues making direct contact with A2ʹ of the G**A**AA tetraloop, A8-U3. Most mutations at this base step resulted in deleterious effects on binding beyond the limit of detection with G**A**AA but several mutations have ΔΔ*G* < 1 kcal/mol with the G**U**AA tetraloop (Fig. 3*B* and *C*, orange). These diminished effects likely arise because the interaction made between the A8-U3 base step and A2ʹ of the G**A**AA tetraloop (the UA-handle) is not present with the G**U**AA tetraloop and thus cannot be further disrupted.

Mutations to the A4-A5 platform that reduced G**A**AA tetraloop binding by 2.0 – 2.5 kcal/mol barely affected binding to the G**U**AA tetraloop (Fig. 3*B* and *C*, red), suggesting that the dinucleotide platform does not significantly contribute to G**U**AA binding. U7-G6 mutations also had diminished effects, consistent with their involvement in positioning the A4-A5 dinucleotide platform, which is not utilized with the G**U**AA tetraloop (Fig. 3*B* and *C*, purple).

All but one mutation to the G11-C1 and G10-C2 decreased binding beyond the limit of detection for binding to G**U**AA (Fig. 3*C*, blue and green). Nevertheless, the mutant with a measurable Δ*G*_bind_ gave a smaller decrease in binding compared to G**A**AA (smaller ΔΔ*G*), suggesting an energetic interaction, presumably indirect, between these base steps and A2ʹ of the tetraloop.

Overall, the A2ʹ→U tetraloop mutation decreased the single mutation effects across the 11ntR, indicating broad energetic coupling to the 2ʹ residue of the tetraloop. While coupling is expected for the residues making direct interactions, the decreased effects on binding for the other 11ntR residues suggest that these single mutants induce conformational penalties for binding.

### Idiosyncratic effects upon base partner substitutions

RNA base pairing typically follows simple rules, namely the formation of Watson Crick, wobble pairs, and sometimes other pairs with specialized geometries and/or interactions (44, 45). Correspondingly, RNA structure and function is often rescuable by restoring base complementarity, and this property has been used extensively to identify helices in complex RNAs (46–48). Cases in which rescue is incomplete have been suggested as evidence for tertiary interactions (*e*.*g*., (49, 50)). Prior studies have typically been limited in the number of mutants that can be studied or in their ability to quantitatively probe thermodynamic behavior, which limits the ability to explore base pair patterns and potentially masks underlying structural and functional properties. We took advantage of the high throughput and quantitative nature of RNA-MaP to thoroughly determine base partner preferences throughout the 11ntR.

In the 11ntR, all but one of the residues (9U) make base-base interactions with a partner residue, and these interactions involve opposing strands of the 11ntR in all cases except for the 4A-5A (Fig. 1*C*). There are two canonical Watson-Crick base pairs (11G-1C and 10G-2C), one wobble pair (7U-6G), and two non-canonical base pairs (4A-5A, 8A-3U). We investigated the effects of all 16 base pair substitutions at each of these five base steps for binding the GAAA tetraloop and uncovered expected and unexpected patterns of effects on Δ*G*_bind_ (Fig. 4A).

**Fig. 4.**
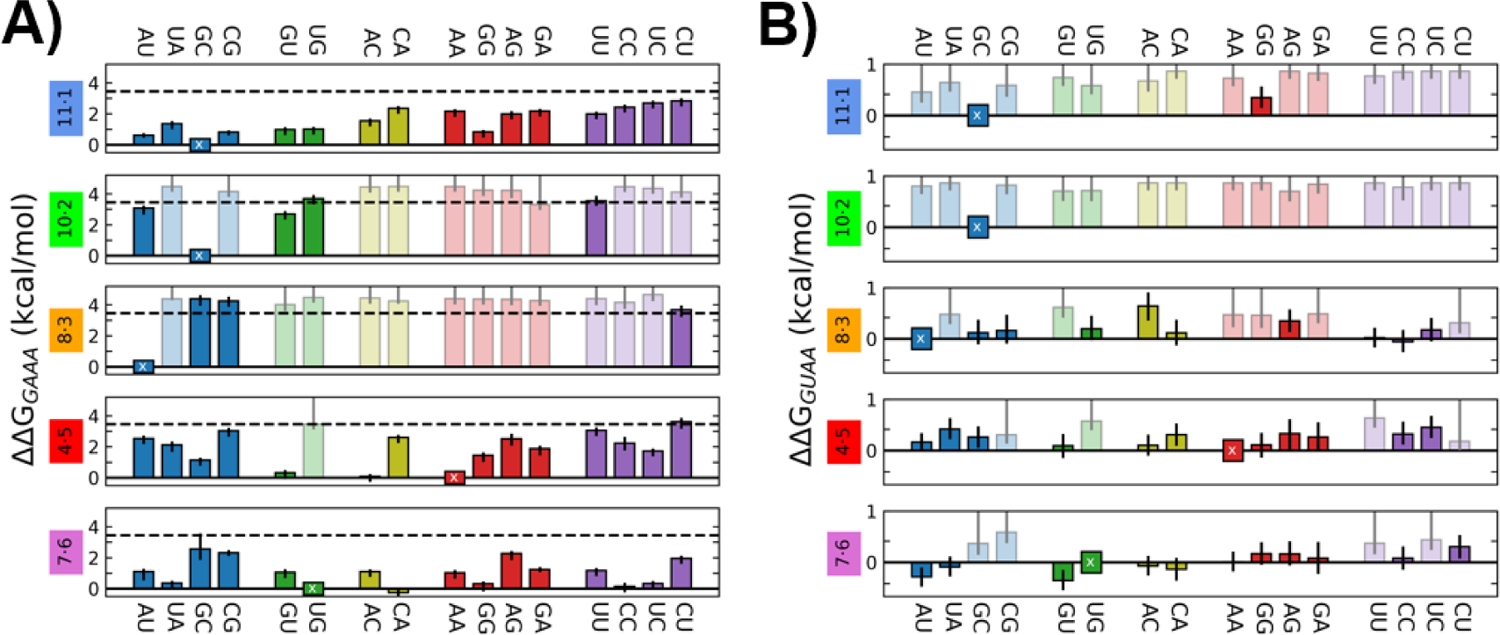
Systematic base partner substitutions to the 11ntR reveal idiosyncratic behaviors. Energetic effects (ΔΔ*G*) of base partner substitutions in the 11ntR for binding to GAAA (A) and GUAA (B) tetraloops. All possible base pair substitutions are shown at each base pair level of the 11ntR, as described in Fig. 3A. The ΔΔ*G* between wild-type 11ntR binding to GAAA *versus* GUAA is presented for reference as the dashed line in (A). The substitutions are grouped by type: Watson-Crick (blue), wobble pair (green), RY mismatch (yellow), RR mismatch (red), and YY mismatch (purple). The wild-type sequence at each base step has ΔΔ*G* = 0 and is denoted with an “x”. Error bars represent 95% confidence intervals for ΔΔ*G* values, and substitutions that result in limits are denoted by translucent boxes.

Starting with the core residues, we found that all substitutions to the base steps 10G-2C and 8A-3U greatly weaken binding, with most substitutions decreasing binding beyond the limit of detection (ΔΔ*G* > 3.5 kcal/mol). Additionally, most of the substitutions that did result in detectable binding had affinities similar to that of the wild-type 11ntR • G**U**AA interaction (dashed line in Fig. 4A), in which one of the core 11ntR • GAAA interactions is removed. This absence of rescue to binding at the core base pairs is expected given their extensive complex interactions with the tetraloop.

The energetic effects of substitutions to step A4-A5, the “dinucleotide platform,” are more varied than those at the core, with two mutations yielding almost no change in Δ*G*_bind_ and others decreasing binding to near and beyond the limit of detection. The substitutions with ΔΔ*G* < 0.5 kcal/mol, G4-U5 and A4-C5, are also found as dinucleotide platforms across different structured RNAs, including other natural 11ntRs (51, 52). The ability to substitute platforms at the A4-A5 base step — relative to the core base pairs involved in tertiary interactions — may reflect the lower specificity of stacking relative to hydrogen bonding. Six of the other substitutions, U4-C5, G4-G5, C4-A5, U4-A5, U4-U5, and C4-G5, form platforms in other RNAs yet exhibit a range of binding energies in the 11ntR • GAAA interaction (ΔΔ*G* = 1–3.5 kcal/mol; Fig. 4A; (51)). These differences presumably reflect, in part, differences in stacking energies and may reflect the interplay of stacking energy with base pair step geometry (53–55). The platform also is energetically connected to the 8A-3U core base pair (see *Energetic cooperativity within the 11ntR is strong, extensive, and incomplete* below) so that platform changes may also influence the conformation of 8A-3U and its ability to form the UA-handle interaction.

Substitutions at the peripheral base steps G11-C1 and U7-G6 affect tetraloop binding despite lacking direct interaction. These effects are generally more modest than those in the core but nevertheless vary considerably with base substitutions. Replacing the wild-type G11-C1 with another Watson-Crick pair (Fig. 4*A*, blue) results in moderate effects on Δ*G*_bind_ of 0.5-1.5 kcal/mol, and wobble pair substitutions result in similar ΔΔ*G* values (Fig. 4*A*, orange). The 11G-1G mismatch (green) also gives a small effect whereas all other mismatches give larger deleterious effects of > 2 kcal/mol (yellow, red, and purple). The U7-G6 substitutions result in even more idiosyncratic behaviors, with some base partners giving almost no effect on and others giving effects of >2 kcal/mol (Fig. 4*A*).

In the simplest scenario, the energetic effects of base step substitutions should follow their geometric similarities, *i*.*e*., isosteric substitutions at base steps would yield similar ΔΔ*G* values. However, we find little to no correlation of the energetic effects from base pair substitutions with the IsoDiscrepancy Index, a metric defined by Stombaugh *et al*. (44) to quantify isostericity between base pairs (*SI Appendix* Fig. S6). The substitution effects at these peripheral base steps suggest that mutating these residues alter the conformational preferences of the core residues that directly participate in tertiary interactions but do so in non-obvious ways. Further, these results indicate that an absence of rescue by a seemingly canonical substitution need not indicate a direct tertiary interaction to that base pair.

Base pair substitutions gave smaller effects on binding the G**U**AA tetraloop relative to the G**A**AA tetraloop (Fig. 4*B*). These results are similar to the single mutation effects and are again consistent with the presence of fewer tertiary contacts — *i*.*e*., less binding energy to lose (Fig. 3*A* and *B*). In particular, core base partner substitutions at U8-A3 that resulted in ΔΔ*G* > 3 kcal/mol with G**A**AA yielded small or negligible effects (ΔΔ*G* < 1 kcal/mol) with G**U**AA. In addition, no U8-A3 substitution significantly increased binding (*i*.*e*., all ΔΔ*G* ≥ 0), providing no indication of an alternative to the native UA-handle • G**A**AA interaction with the G**U**AA tetraloop. Finally, some substitutions at U7-G6 provide modest benefits for G**U**AA binding (ΔΔ*G* < 0, *SI* Appendix Table S1), potentially reflecting small changes in the conformation of the receptor. As of yet, there are no structures available for the 11ntR bound to the G**U**AA, and structural characterization of these mutants will aid in interpreting our observed energetic effects.

### Energetic connectivity throughout the 11ntR

We investigated the energetic interconnectivity throughout the 11ntR via double mutant cycles for all positions within the 11ntR. Cooperativity is generally viewed as the hallmark of tertiary structure, and there have been numerous discussions of cooperative behavior in RNA folding (50, 56–58). Our high-throughput approach allowed us to systematically look for cooperativity between all residues of the 11ntR and, more generally, to ascertain the nature of the energetic connectivity throughout the 11ntR, as defined in Fig. 5*A* and *B*.

**Fig. 5.**
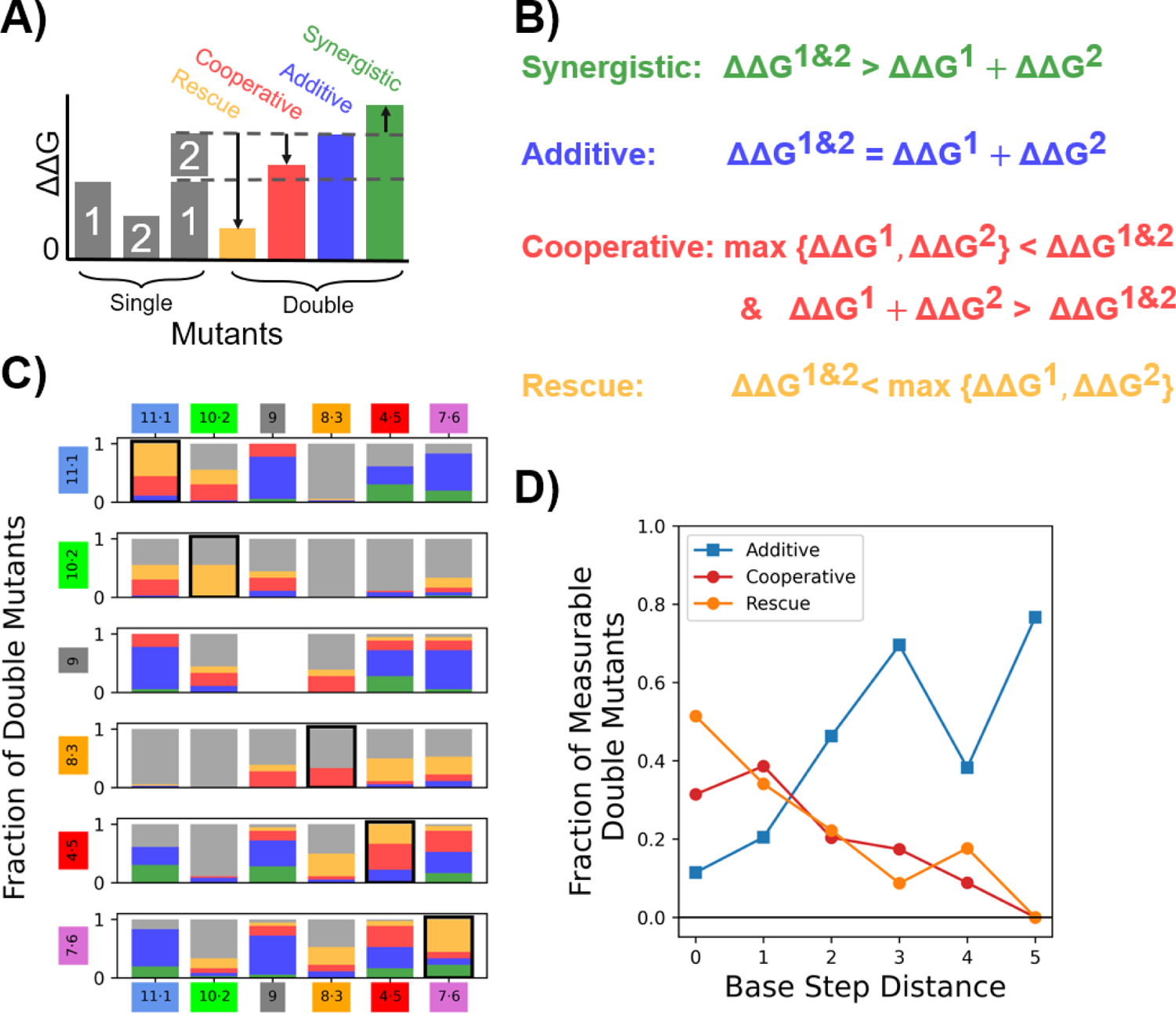
Energetic coupling within the 11ntR defined by double mutant cycles. Energetic effects from double mutant cycles can be classified by comparing the double mutant ΔΔ*G* value to its constituent single mutant ΔΔ*G* values. The classes are rescue, cooperative, additive, and synergistic as depicted schematically in (A) and by the equations in (B). (C) Energetic coupling classes grouped by base pair steps; classes are color-coded as in (A) and (B), with gray denoting unmeasurable coupling due to one or more measurements being a limit. Each stack represents nine residue combinations within each base step for boxed elements along the diagonal, 36 combinations for interactions between base steps, or 18 combinations for interactions of each base step with residue U9. (D) The fraction of additive, cooperative, and rescue mutants — out of the total of measurable (non-limit) mutants — plotted against the base step distance between the two mutations.

Fig. 5*C* summarizes the types of energetic connectivity observed for the double mutants, grouped by base step. The multiple types of energetics for most pairs of positions (multi-color bars in Fig. 5*C*) indicate that the base identity of mutations affect the energetic signature, revealing an additional level of complexity to energetic relationships. For example, base-specific effects at G11-C1 result in rescue when a double mutant is a Watson-Crick pair but result in cooperativity otherwise (*SI Appendix* Fig. S7). Likewise, we observe rescue and cooperativity within base pair steps (black outlines), and these connections extend to their nearest neighbors. Connectivity is expected for bases directly interacting across from one another, and the energetic connection between neighboring base partners echoes the “nearest neighbor rules” where the identity of a base pair and only its nearest neighbors are required to accurately predict duplex stability and conformational ensemble properties (38, 59, 60). Even U9, which flips out in the tertiary structure, exhibits cooperative energetics with its neighbors (the G10-C2 and U8-A3 steps). Additive energetics (Fig. 5*C*, blue) are observed throughout the motif, but more frequently in residues further from one another.

Overall, energetic coupling between residues decreases with distance along the motif, as seen in Fig. 5*D*. The number of cooperative and rescue (red and orange) mutants, indicative of energetic coupling, decreases with separation, whereas additive mutants (blue) increase with separation. This gradient in connectivity — rather than a sharp drop-off — indicates that the 11ntR is not comprised of energetically independent sub-regions. Thus, while submotifs, such as the A-minor interactions and the A–A platform, have been identified across many RNAs, this ability to distinctly visualize these ‘motifs’ within RNA structures does not necessarily indicate that they act as energetically distinct units. Our dissection of the energetic architecture of the 11ntR paints a more nuanced picture of the motif. For example, we find that the G11-C1 base step alters the energetics of “A-minor” motif formed by the G10-C2; the A4-A5 and U7-G6 base steps alter the “UA-handle” formed by U8-A3; and the U7-G6 base step alters the “A–A platform” formed by A4-A5. But despite these connections, cooperativity within the 11ntR is far from complete, a property that leads to a more complex energy landscape that likely aids the stepwise evolution of RNA-RNA interactions.

### Synergetic energetic penalties to 11ntR • GAAA formation

Synergistic effects make up the smallest fraction of energetic connectivity and is the least intuitive type of energetic interaction, as it entails a second mutation having a larger unfavorable impact in the presence of a first mutation (Fig. 5*A* and *B*). Synergistic effects can arise from so-called “threshold effects”, where the energetic effect (from two mutations together) arises from unfolding or formation of another type of non-optimal state such as nonfunctional secondary structures that become the dominant state only when both mutations are present (*e*.*g*., (61, 62)). For the 11ntR, the A–A platform appears to be especially sensitive to synergistic effects, with double mutants involving residues A4, A5, G6, and U7 resulting in the largest synergistic effects (Fig. 5*C*, *SI Appendix* Fig. S7). Many of these double mutants appear to result in alternative secondary structures (*SI Appendix* Table S2).

We propose two specific models that can account for these effects. First, double mutations that favor base pairing interactions between A4 or A5 and the opposing strands lead to synergy, as the constituent single mutants are not as prone to the formation of such structures. Second, double mutants that lead to the formation of a Watson-Crick base pair at U7-G6 (which is deleterious based on single mutations that form such base pairs; Fig. 3*B*) give rise to synergy. The exception is a U7-A6 mutant, which has previously been proposed to form a non-Watson-Crick base pair with a geometry more akin to a wobble pair (33). Thus, dinucleotide platforms and their neighbors may be subject to increased evolutionary pressures, as changes to their sequence or local environments can lead to alternative structures and decreased function.

### The conformation of the 11ntR • GNAA is robust to mutation

While the extensive energetic effects and energetic interrelationships strongly imply coupled conformational effects within the 11ntR *•* GAAA, the orientations of the helices that emanate from this motif appear to be unchanged based on the observation of indistinguishable thermodynamic fingerprints for measurable mutants (*SI Appendix* Fig. S5*A*; (36, 40)). The five scaffolds studied provide a range of approach orientations for the 11ntR • GAAA interaction, and the mutants follow the same affinity pattern as wild-type across the scaffolds, despite weaker overall binding.

These results extend to GUAA binding. The fingerprints for the wild-type 11ntR *•* GAAA and wild-type 11ntR *•* GUAA are not significantly different (*SI Appendix* Fig. S5*C* and *D*), and no single mutant results in a greater specificity to the alternative tetraloop (ΔΔ*G*_GUAA_ ≥ ΔΔ*G*_GAAA_ for single mutants). Further, only one single mutant, 8A→G, has a significantly different fingerprint compared to the wild-type 11ntR *•* GAAA interaction (*SI* Dataset S3), so the receptor generally maintains its overall “motif geometry” despite changes to the local conformation of the receptor. These results indicate that the sequence and conformational specificity of the 11ntR *•* GAAA interaction extends to the immediate sequence space around the wild-type interaction (two total mutations to the receptor and/or tetraloop). Nevertheless, four double mutations increased binding to the GUAA tetraloop (*SI Appendix* Table S1), and 26 double mutants showed evidence of scaffold effects (*SI Appendix* Fig. S5*B* and *E*). Thus, the tertiary contact diverges from 11ntR *•* GAAA conformational behavior for a small subset of variants with three mutations: two to the receptor and one to the tetraloop.

### Exploring evolutionary driving forces for the 11ntR beyond tertiary contact strength

Bonilla *et al*. (40) previously found that the binding energy of 70 11ntR variants observed in nature correlate with their abundance in RNAs (*ρ* = −0.47). As we investigated all single and double 11ntR mutants (528 + the wild-type 11ntR), we could extend this analysis to include many variants not observed in nature and test the prediction that 11ntR sequences not observed in nature form less stable tertiary interactions than those that are found in nature. These variants not found in nature have likely been selected against during the evolution of the 11ntR and thus could yield insight into the evolutionary constraints on the tertiary contact. Indeed, 88% (434 of the 496 single and double mutants not observed) were less stable than the least stable natural 11ntR variant, consistent with the prior conclusion that motif stability is an important constraint (Fig. 6*A*).

**Fig. 6.**
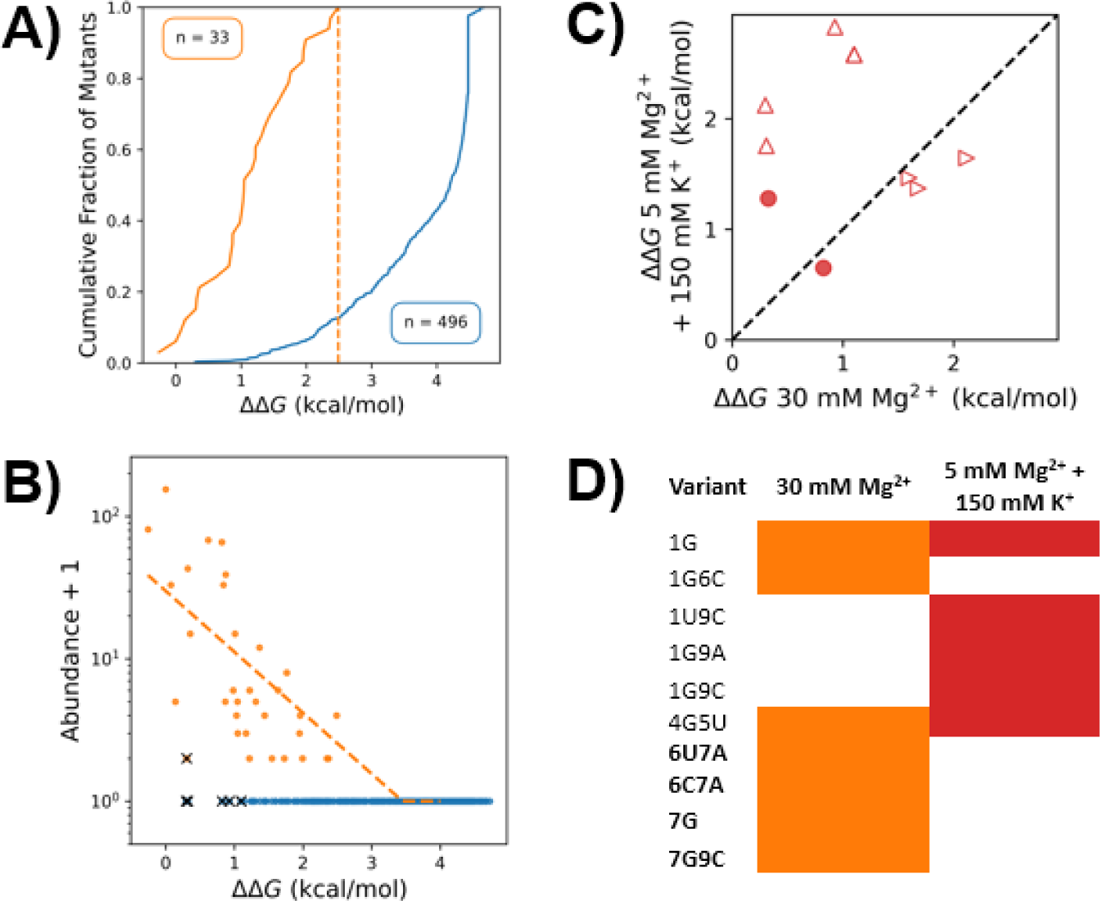
The abundance of 11ntR variants correlate with their stability relative to wild-type. (A) The cumulative fraction of natural 11ntR variants is plotted against their stability in 30 mM Mg^2+^ as an orange line; the cumulative fraction of non-observed variants is shown as a blue line. Dashed lines represent the ΔΔ*G* value of the least stable natural variant. (B) There is good correlation between stability and abundance for the natural variants (orange) under 30 mM Mg^2+^, as noted in Bonilla *et al*. (40); variants not observed in nature are in blue. The dashed line represents an exponential fit, described in the *Methods*; black “x”s represent variants with high stabilities (low ΔΔ*G* values) that have abundances in natural RNAs that are significantly lower than expected given the correlations (*SI Appendix* Table S3). (C) Stabilities of significantly underrepresented abundances in either 30 mM Mg^2+^ or 5 mM Mg^2+^ + 150 mM K^+^; variants that are significant in both conditions are denoted by a filled circle; variants significant only in 30 mM Mg^2+^ are denoted by ►; variants significant only in 5 mM Mg^2+^ + 150 mM K^+^ are denoted by ▲. (D) The outlier mutants as identified in (C) color coded as orange for 30 mM Mg^2+^ and red for 5 mM Mg^2+^ + 150 mM K^+^. Mutations to the K^+^ binding site are bolded.

Of the single and double 11ntR mutants, seven variants had abundances significantly lower than expected based on stability alone (Fig. 6*B*, denoted by “x”). The simplest model to account for these discrepancies is the non-physiological assay conditions used in RNA-MaP (30 mM Mg^2+^ used to enhance the assay’s dynamic range; (36)). Indeed, Bonilla *et al*. (40) found that the correlation increased for the 70 11ntR variants observed in nature under more biologically relevant conditions, with lower Mg^2+^ and K^+^ present (*ρ* = −0.47 *vs*. −0.69). Under these conditions (5 mM Mg^2+^ & 150 mM K^+^), the abundances of four of the seven outliers were no longer lower than expected (Fig. 6*C*;*SI Appendix* Fig S8) and all four mutated the site of K^+^ binding, the U7-G6 base step (Fig. 6*D*; (27, 29, 63–65)).

In contrast, the G4-U5 substitution remains an outlier in the presence of K^+^ (Fig. 6*D*). While we found this substitution to be energetically equivalent to the native A4-A5, much less frequently in native 11ntR-containing RNAs (Fig. 4*A*; (24, 25)). Geary *et al*. (33) proposed that this underrepresentation could result from alternative secondary structures formed *in vivo* that do not arise under the experimental conditions. Other significant outliers (present less than predicted) arise from mutations at position 1 or positions 1 and 9 but lack obvious potential origins. Overall, these results further highlight the power of systematic and quantitative investigation of sequence space to identify outliers for further studies that may reveal new structural, energetic, and functional behavior.

## Conclusions and Implications

Our quantitative analysis of all single and double 11ntR mutants binding to the canonical GAAA tetraloop and the suboptimal GUAA tetraloop provide an unprecedented energetic dissection of this RNA motif. Comprehensive double mutant cycle analysis allowed us to dissect the energetic architecture of the 11ntR. Whereas RNA motifs and structures have widely been described as highly cooperative, we uncovered a more nuanced energetic architecture: base partners and stacking neighbors commonly exhibit cooperativity and sometimes rescue, indicative of intimate conformational and energetic interrelationships. We also obtained evidence of long-range cooperativity between peripheral residues and the interaction core, indicating energetic linkage via alterations to the local conformational ensemble. Nevertheless, these energetic interactions occur as a gradient, falling off with distance across the motif, rather than exhibiting complete cooperativity within the motif or energetically independent sub-motifs. Structural units that have been described as motifs, such as the A–A platform or A–minor motifs, may not behave as energetic units within the 11ntR, as suggested from cooperativity between these so-called motifs and the rest of the motif. Conversely, the 11ntR motif as a whole appears to behave as a transferable energetic unit across sequence contexts and environmental conditions, in accordance with the Reconstitution Model of RNA folding (20).

Base partner substitutions within the 11ntR recovered expected behaviors, namely Watson-Crick rescue within base pairs that do not interact with the tetraloop and limited rescue for substitutions at sites that do make direct tertiary interactions. Nevertheless, unexpected base partner preferences were also observed, even at sites within the motif that do not make direct interactions. These idiosyncrasies underscore a more complex energetic architecture of the 11ntR motif relative to simple helical RNA segments, wherein isosteric substitutions generally give energetic and conformational equivalence (38, 44, 45). These observations provide targets for future RNA structural studies to uncover the origins of the near-native conformational and energetic behavior of particular non-canonical base pair substitutions to the 11ntR.

With RNA-MaP, we are able to map sequence changes to energetic consequences throughout RNA motifs. The thorough coverage of sequence space (of all single and double mutants) allowed identification of sequences that gave both unexpectedly stronger and weaker tertiary binding than anticipated by base partner identities or additivity. Further, the sequence space explored by such a high-throughput technique enables the study of both natural and non-natural variants of a motif. Such studies are pivotal in defining the energetic landscape traversed through evolution and ultimately in understanding how sequence and structure relate to energetics and nature’s choices throughout evolution.

## Methods

### Data Curation

Given their large scale, RNA-MaP experiments are typically designed to simultaneously address multiple questions. The data used in this study is a subset of the data broadly presented in Bonilla *et. al*. (40). New binding isotherms were fit to the normalized fluorescence data as previously described to facilitate the generation of larger bootstrap sample sizes (*n* = 10,000; (36, 38, 40)). In short, empirical bootstrapped distributions for binding free energies were derived from resampling the normalized fluorescence data for molecular replicates (clusters) and fitting binding isotherms to Equation 1:

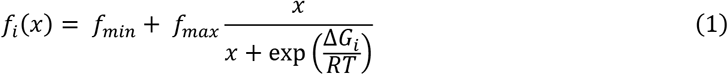

where *x* represents flow piece concentration (8 total concentrations ranging from 0.9 to 2000 nM), *f*_*i*_(*x*) is the resampled normalized fluorescence for a given flow piece concentration, *R* is the ideal gas constant, *T* is temperature in Kelvin, *f*_*min*_ and *f*_*max*_ are parameters that describe the minimum and maximum normalized fluorescence across all clusters, and Δ*G*_*i*_ is the fit free energy for the resampled fluorescence. We use the index *i* to reference bootstrap replicates herein.

We set a minimum number of five clusters per sequence variant to fit equilibrium constants. Sequence variants had an average of 65 and 32 clusters for binding to GAAA and GUAA, respectively (*SI Appendix* Fig. S2*A* and *B*). Two experiments were performed to measure binding of 11ntR variants to the GAAA tetraloop; we combined data from the experimental replicates by averaging the two Δ*G*_bind_ values for each bootstrap replicate by Equation 2:

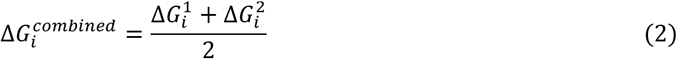

where 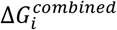 is the averaged Δ*G*_bind_ value and 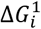 and 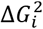 are the Δ*G*_bind_ values from each replicate.

Δ*G*_bind_ values for the 5 mM Mg^2+^ + 150 mM K^+^ condition were used with the original error estimates from Bonilla *et al*. (40) because they were not used to perform additional statistical tests herein.

Due to the experimental threshold for binding of –7.1 kcal/mol, library variants with a median Δ*G*_bind_ ≥ –7.1 kcal/mol had their Δ*G*_bind_ set to –7.1 kcal/mol, which is a lower limit for their Δ*G*_bind_ values.

### Statistical analysis

We leveraged the empirical distributions of Δ*G*_bind_ obtained via bootstrapping to directly propagate uncertainty throughout our calculations. Reported Δ*G*_bind_ values and ranges in the text represent the median and 95% confidence interval of the distribution from the average of Δ*G*_bind_ values over the five scaffolds by Equation 3:

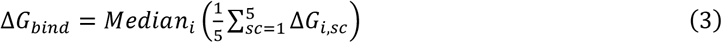

where Δ*Gi,sc* represents the bootstrapped replicates for Δ*G*_bind_ indexed by *i* for each of the five tectoRNA scaffolds indexed by *sc* and the median is taken over the bootstrap replicates. ΔΔ*G* values, the difference between variant and wild-type binding affinities, were calculated by subtracting the wild-type Δ*G*_bind_ value from the mutant Δ*G*_bind_ value for each scaffold and averaging over the scaffolds. Reported ΔΔ*G* values in the text reflect the median and confidence interval from the bootstrapped distributions, by Equation 4:

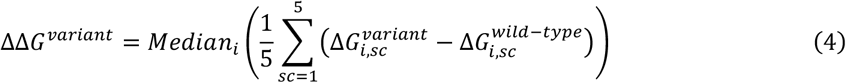

Energetic coupling analysis was performed by sequentially testing the null hypotheses that there is no synergistic effect, no rescue effect, and no cooperative effect (as defined in Figure 5*A* and *B*); *p*-values for these hypotheses were calculated by Equations 5-7:

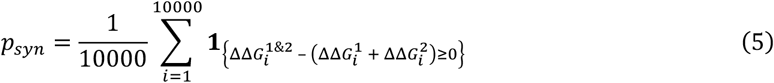

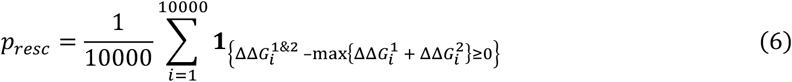

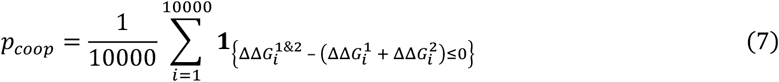

where 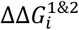 are the bootstrapped ΔΔ*G* values of a double mutant; 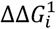 and 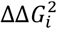 are the bootstrapped ΔΔ*G* values for its constituent single mutants; **1**{•} is the indicator function (represented by 1 if the condition holds and 0 if it does not); and *p*_*syn*_, *p*_*resc*_, and *p*_*coop*_ are the probabilities (*p-*values) that there is no synergistic effect, no rescue effect, and no cooperative effect, respectively. These *p-*values were corrected using Benjamini*–*Hochberg false discovery rate correction for all 1485 tests. Double mutants were classified as synergistic if *p*_*syn*_< 0.05 and rescue if *p*_*resc*_ < 0.05; unclassified mutants with *p*_*coop*_ < 0.05 were classified as cooperative since rescue is a stronger form of cooperativity; the remaining unclassified mutants were classified as additive, which represents the condition given in Equation 8:

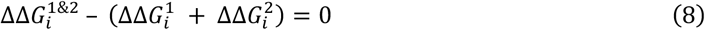

Relative free energy values, *p*_*syn*_, *p*_*resc*_, and *p*_*coop*_ for all 495 11ntR double mutants binding to the GAAA or GUAA tetraloop are reported in *SI* Dataset S2.

We identified changes to the binding conformation of 11ntR mutants (i.e., “thermodynamic fingerprints”) by testing the null hypothesis that a 11ntR variant and the wild-type 11ntR had the same fingerprints, *i*.*e*., test if the ΔΔ*G* values zero variance over the five scaffolds, within error (Figure 2*C*). We first estimated the expected distribution of variance in the ΔΔ*G* values from experimental error by computing the variance in ΔΔ*G* values between the wild-type 11ntR • GAAA Δ*G*_bind_ values and a permuted set of wild-type 11ntR • GAAA Δ*G*_bind_ values, *i*.*e*., the variance in ΔΔ*G* values of the wild-type and itself. This permutation was done by shifting the bootstrap index, *i*, of the wild-type Δ*G*_bind_ replicate values by one as by Equation 9:

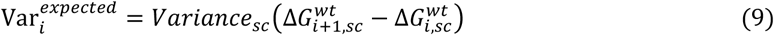

where the variance for each bootstrap replicate *i* is calculated over the scaffolds *sc*.

Since the null model is that there is no variance in ΔΔ*G* values, we used Equation 10 to calculate the probability (*p*-value) that the variance is within the expected range for error (based on Equation 9) for each 11ntR variant (for binding to the GAAA or GUAA tetraloop):

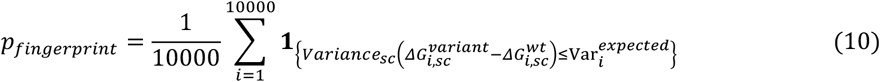

where again, the variance for each bootstrapped replicate *i* is calculated over the scaffolds *sc* and is compared to the bootstrapped expected variance 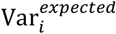.

We then applied Benjamini*–*Hochberg false discovery rate correction to correct the *p*-values for multiple hypotheses testing. We rejected the null for *p*_*fingerprint*_ < 0.05 and concluded that these variants had different fingerprints *versus* the wild-type 11ntR • GAAA interaction. The median and 95% confidence intervals for the variance in ΔΔ*G* values of 11ntR variants binding to the GAAA or GUAA tetraloop relative to the wild-type 11ntR • GAAA interaction and the resulting *p*_*fingerprint*_ values are listed in *SI* Dataset S3 for variants with data for 3 or more scaffolds (293/528 for GAAA binding and 238/529 for GUAA binding).

### Identifying unexpectedly low abundance 11ntR variants

We analyzed a subset of natural 11ntR variants presented in Bonilla *et al*. (40), which established an exponential correlation between abundance and stability. This correlation makes predictions for the abundance of variants based on their stabilities, including variants that are not observed in nature. To recapitulate this correlation and identify stable variants with significantly lower abundances than predicted by their stability, we fit an exponential relation between [1 + Abundance] and stability (ΔΔ*G*) to 33 variants of the 70 natural variants identified in Bonilla *et al*. that were either wild-type, single, or double mutants. As abundance cannot be negative, we enforced a minimum of zero abundance such that the final model is given by Equation 11, where *β*_*0*_ and *β*_*1*_ are free parameters for the fit.

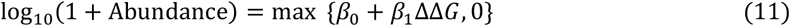

We then calculated residuals between the predicted abundance and actual abundance of all 529 11ntR variants (with mutants not found in nature having zero actual abundance) using Equation 12. Negative residuals correspond to variants whose abundances are lower than expected given their stabilities; positive residuals represent variants whose abundances are higher than expected.

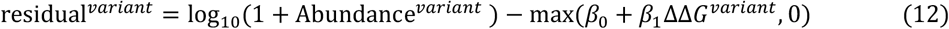

To identify variants that are statistically significantly under and overrepresented in nature based on their stability, we computed the z-scores for all residuals, identifying seven variants with z-scores below a Bonferroni-corrected *p*-value of 0.05 and one variant with z-scores above the threshold (Equations 13 and 14, where *Φ*^−*1*^ represents the quantile function for the standard normal distribution).

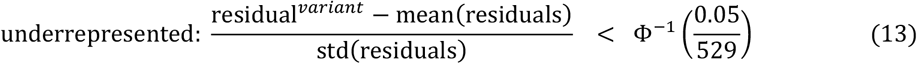

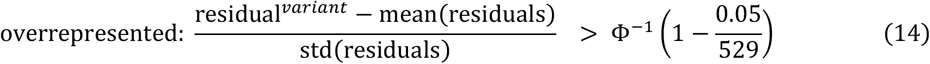

We repeated this procedure using ΔΔ*G* values obtained under a different salt condition (5 mM Mg^2+^ + 150 mM K^+^). Three additional three variants with significantly lower abundances and one variant with significantly higher abundance were identified, relative to the 30 mM Mg^2+^ conditions. The variants that differed significantly and their ΔΔ*G* values and z-scores are listed in *SI Appendix* Table S3.

## Data, Materials, and Software Availability

All study data are included in the article and/or supporting information. Code is available upon request.

## Acknowledgments

We thank members of the Herschlag laboratory for discussion and review of the manuscript. This work was supported by the National Institutes of Health (R01 GM132899 to D.H.). J.H.S. was supported by the NSF Graduate Research Fellowship Program under Grant DGE-1656518.

