## Supplemental_Materials for "Dissecting the energetic architecture within an RNA tertiary structural motif via high-throughput thermodynamic measurements": Shin-11ntR_RXiv_Supplement.pdf

```
Scaffold 1   Scaffold 2   Scaffold 3   Scaffold 4   Scaffold 5

                                         g-a
                                         g a          g-a
   g-a          g-a          g-a         g-c          g a
   g a          g a          g a         c-g          g-c
   g-c          g-c          g-c         u-a          c-g
   c-g          g-c          u-a         g-c          u-a
   u-a          a-u          u-a         g-c          g-c
   g-c          c-g          c-g         | g          g-c
   a-u          a-u          u-a         | u          | a
   g g          c-g          a-u         c-g          c-g
   u-a          g-c          g-c         c-g          c-g
   c-g          u-a          g-c         g-c          g-c
   a-u          a-u          g-c         a-u          a-u
   a-u          a-u          a-u         a-u          a-u
   G-C          G-C          G-C         G-C          G-C
   G-C          G-C          G-C         G-C          G-C
   U |          U |          U |         U |          U |
   A·U          A·U          A·U         A·U          A·U
   | A          | A          | A         | A          | A
   | A          | A          | A         | A          | A
   U·G          U·G          U·G         U·G          U·G
   a-u          a-u          a-u         a-u          a-u
   g-c          g-c          g-c         g-c          g-c
   g-c          g-c          g-c         g-c          g-c
   a-u          a-u          a-u         a-u          a-u
   u-a          u-a          u-a         u-a          u-a
   c-g          c-g          c-g         c-g          c-g
```

**Fig. S1.** The five scaffold sequences used in this study, shown with the wild type 11ntR (in orange).

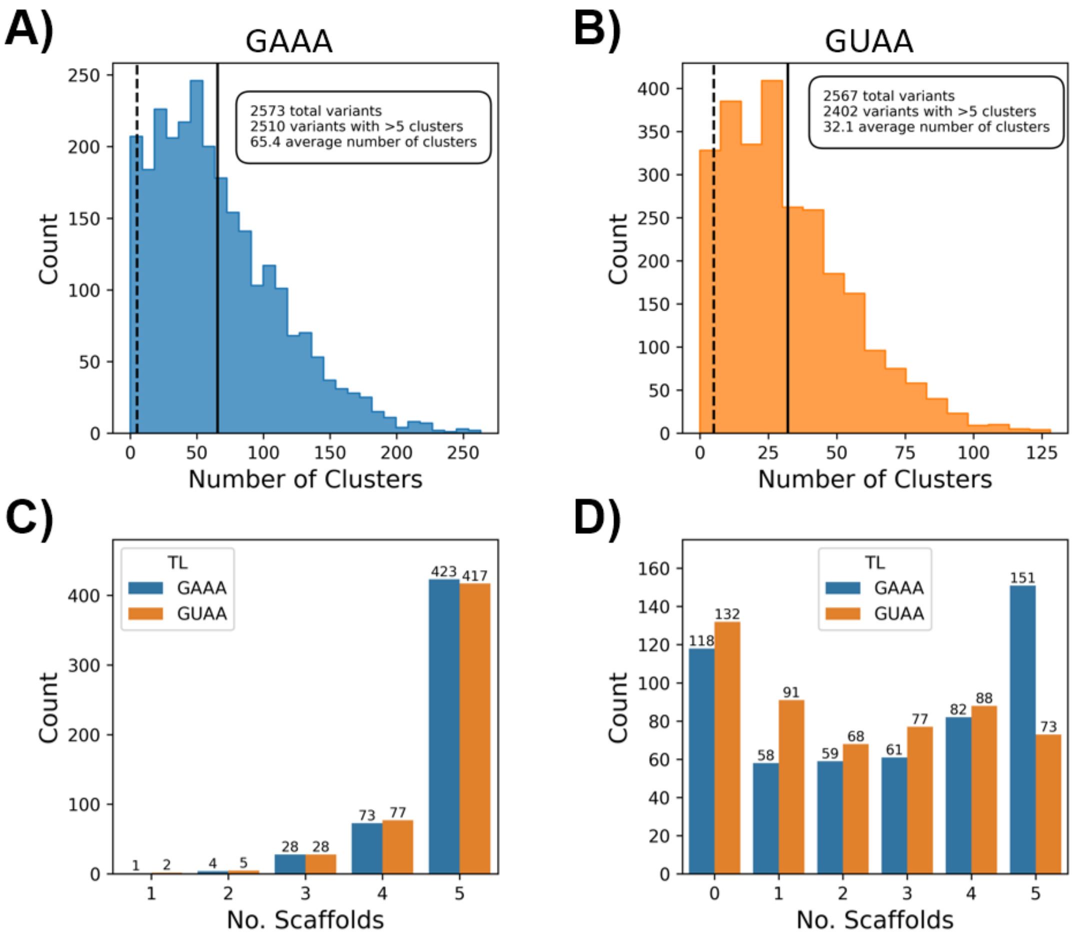


**Figure S2.** The coverage and metrics for 11ntR variants analyzed in this study. The total number of clusters for a given sequence binding to GAAA (A) and GUAA (B). Total variants indicate the number of 11ntR mutant - scaffold combinations (maximum of 2645 combinations for 529 mutants and five scaffolds) that were sequenced on the chip. Dashed lines show the threshold of 5 clusters per variant used in this and prior studies (1–3), and solid lines represent the mean number of clusters. (C) The number of scaffolds (of the five total scaffolds) with binding data to GAAA (blue) and GUAA (orange) per variant. There were 423 and 413 out of 529 total 11ntR variants with all five scaffolds represented in the binding data to the GAAA (blue) and GUAA (orange) tetraloops, respectively. (D) The number of scaffolds with quantitative binding measurements for each 11ntR variant (*i.e.*, not limits; $\Delta G_{bind}$ < -7.1 kcal/mol) for binding to GAAA (blue) and GUAA (orange). Variants with quantitative binding measurements for fewer than five scaffolds are considered limits when averaged in downstream analyses as one or more values used to obtain the average is the limiting value of $\Delta G_{bind}$ = -7.1 kcal/mol (see *Methods* in the main text).

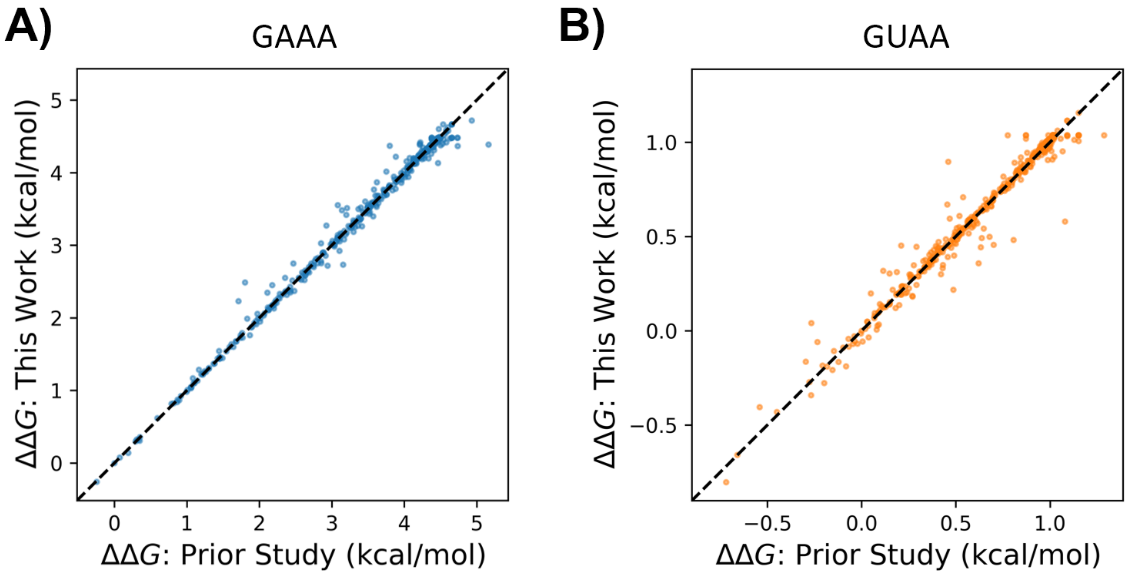


**Figure S3.** Comparison of ΔΔ*G* values from Bonilla *et al.* (3) and after refitting procedure with n = 10,000 bootstrap replicates for binding data to GAAA (A) and GUAA (B) tetraloops. There is overall excellent correlation between fitting procedures, RMSD = 0.09 and 0.06 kcal/mol for GAAA and GUAA binding, respectively. Prior ΔΔ*G* values were within the 95% confidence intervals from refitting for all but two variants: 4C5C for GAAA binding, whose prior and refit ΔΔ*G* are 1.71 and 2.23 (1.77, 2.64) kcal/mol, and 4G8G for GUAA binding, whose prior and refit ΔΔ*G* are 0.81 and 0.48 (0.23, 0.75) kcal/mol.

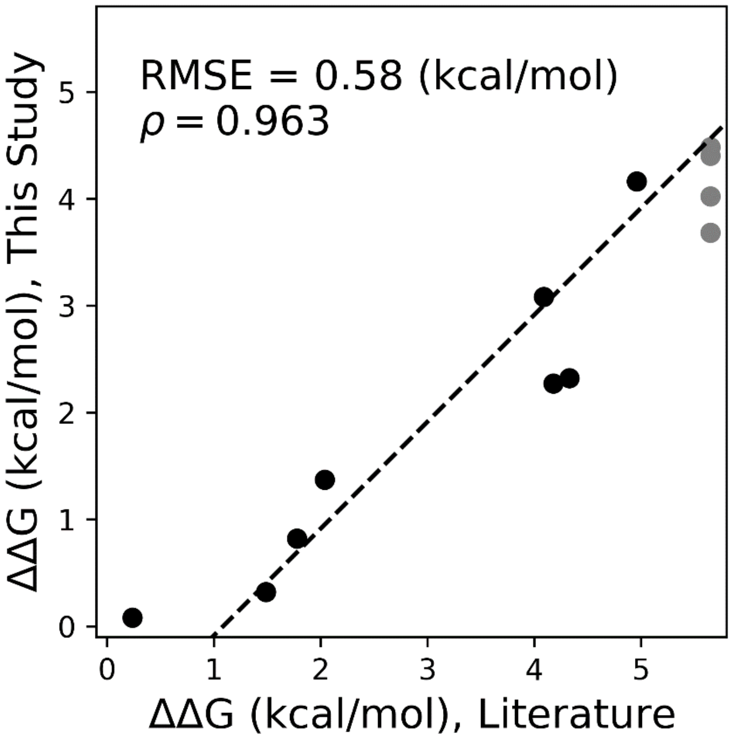


**Figure S4.** Correlation between the ΔΔ*G* values of 11ntR mutants from this study and those from Geary *et al.* (4). Grey points refer to mutants whose binding was at the limit of detection or not measurable in the prior work. ΔΔ*G* values measured in this study were averaged over the five scaffolds denoted in *SI Appendix* Fig. S1 and binding was performed at 22 °C in 30 mM $MgCl_2$; ΔΔ*G* measured in the external study was performed with a different tectoRNA construct at 10 °C and 15 mM $Mg(OAc)_2$.

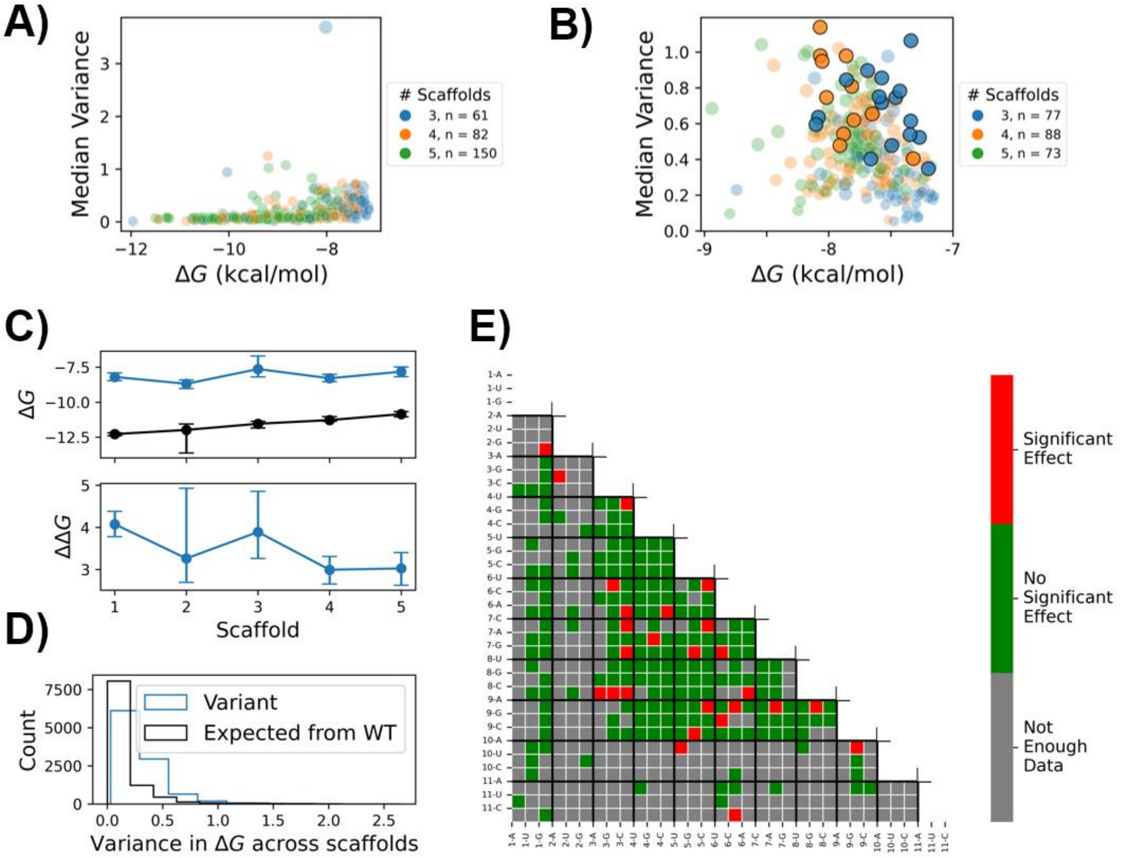


**Figure S5.** Variance in the Δ*ΔG* values between mutant 11ntR variants binding to GAAA (A) or GUAA (B) tetraloops and the wild type 11ntR binding to the GAAA tetraloop. Median bootstrapped variances (described in the *Methods*) are plotted against the average $\Delta G_{bind}$ over the scaffolds. The color represents the number of non-limit scaffolds per 11ntR variant used to calculate the correlations, out of a minimum of three scaffolds and a maximum of five. The size of each point is inversely related to the *p*-value of the null hypothesis that the variance is small (described in the *Methods*). Bigger sizes reflect a higher likelihood of the alternative hypothesis, that there is large variance in the ΔΔ*G* values, *i.e.*, the fingerprints differ. Statistically significant rejections of the null represented are denoted by black outlines. (A) Out of the 293 variants with sufficient binding data to the GAAA tetraloop, no mutant has a significant change in fingerprints. (B) Out of the 238 variants with sufficient binding data to the GUAA tetraloop, one single mutant, 8A→G, and 26 double mutants result in adjusted *p* values < 0.05. (C) The thermodynamic fingerprints (top) for wild-type 11ntR binding to GAAA (black) and GUAA (blue) tetraloops along with the resulting ΔΔ*G* values (bottom) across the scaffolds. The error bars represent 95% confidence intervals based on the bootstrapped distribution for both Δ*G* and ΔΔ*G* values. (D) The distribution of variance in ΔΔ*G* values for the fingerprints in (C) (blue) as well as the expected distribution of variance in ΔΔ*G* values given experimental variability (black). The ΔΔ*G* values between the wild-type 11ntR • GUAA and wild-type 11ntR • GAAA fingerprints were within the expected range, resulting in an adjusted *p*-value of 0.16 > 0.05; thus, the fingerprints are not significantly different. (E) Heatmap indicating which 11ntR double mutants had with significantly different fingerprints when binding to the GUAA tetraloop *versus* the wild type 11ntR • GAAA fingerprint; variances and *p*-values are listed in *SI* Dataset S3.

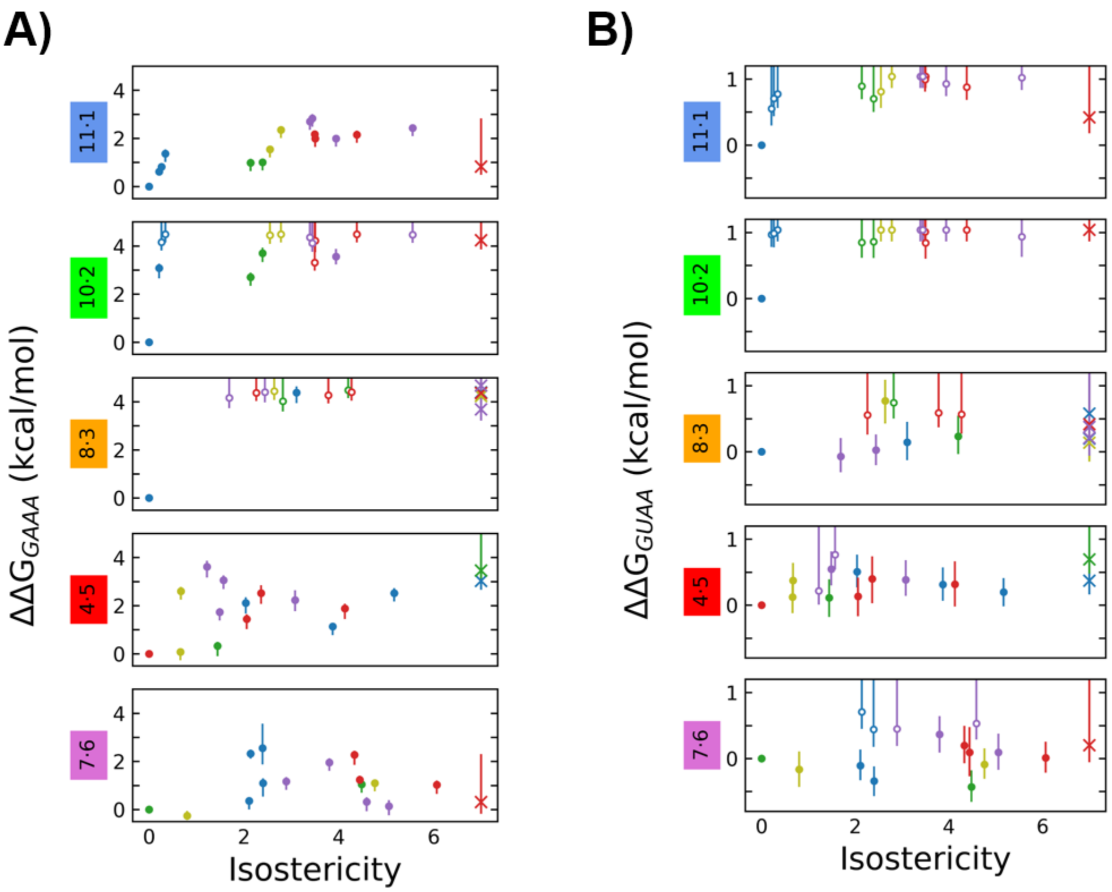


**Figure S6.** Stability *versus* isostericity (quantified by IsoDiscrepancy Index (5)) for base step substitutions to the 11ntR binding to the GAAA (A) and GUAA (B) tetraloops. Substitutions are color-coded by type: Watson-Crick pair in blue, GU wobble pair in green, AC mismatch in yellow, RR mismatch in red, and YY mismatch in purple. Error bars represent 95% confidence intervals in the ΔΔ*G* values, and unfilled points represent limits. Substitutions with no IsoDiscrepancy Index are set to a maximum value of 7 and denoted by “x” (*i.e.*, these base pairs do not exist in RNA structural databases (5)).

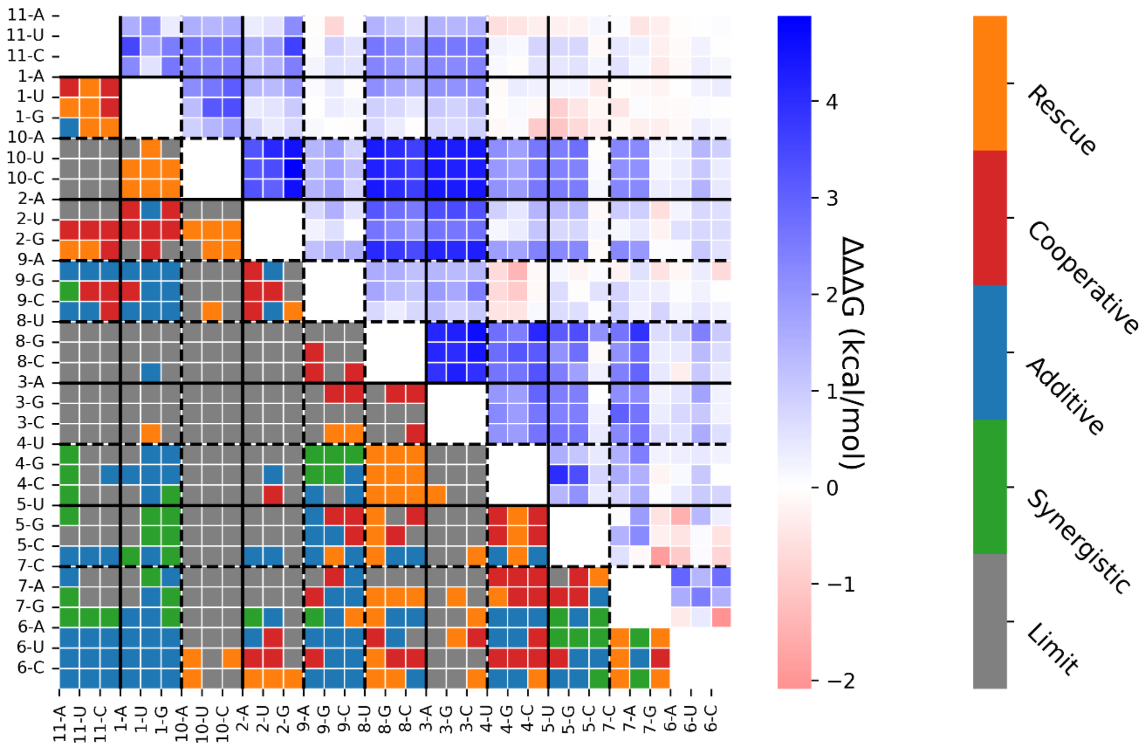


**Figure S7.** Energetic connectivity type and effect size for each double mutant. The lower half of the plot depicts mutants' connectivity types as defined in *Energetic connectivity throughout the 11ntR*. The upper half depicts mutants' effect sizes quantified as follows: $\Delta\Delta\Delta G = \Delta\Delta G^{1\&2} - (\Delta\Delta G^{1} + \Delta\Delta G^{2})$, where cooperative and rescue mutants have $\Delta\Delta\Delta G > 0$, additive mutants have $\Delta\Delta\Delta G \sim 0$, and synergistic mutants have $\Delta\Delta\Delta G < 0$.

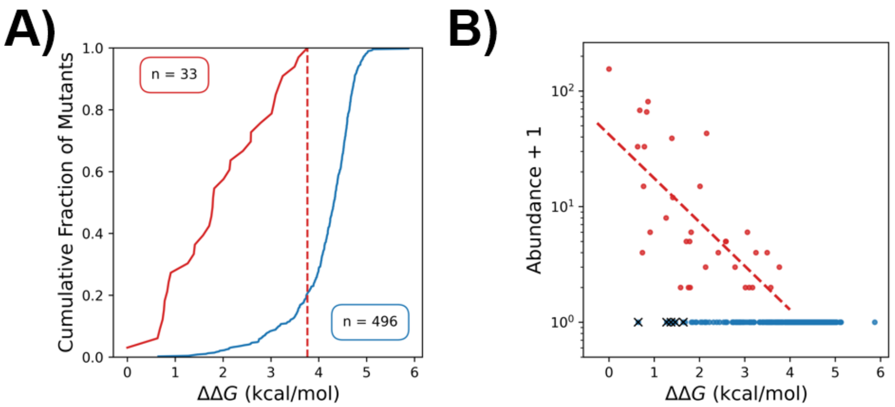


**Figure S8.** The abundance of 11ntR variants correlate with their stability in 5 mM $Mg^{2+}$ + 150 mM $K^{+}$ relative to wild-type. (A) The cumulative fraction of natural 11ntR variants is plotted against their stability in 5 mM $Mg^{2+}$ + 150 mM $K^{+}$ as a red line; the cumulative fraction of non-observed variants is shown as a blue line. The dashed line represents the $\Delta\Delta G$ value of the least stable natural variant. (B) The stability and abundance of natural 11ntR variants (red) correlate well under 5 mM $Mg^{2+}$ + 150 mM $K^{+}$; variants not observed in nature are in blue. The dashed lines are exponential fits, described in the Methods; black "x"s represent variants with high stabilities (low ΔΔG values) that have abundances in natural RNAs that are significantly lower than expected given the correlations (*SI Appendix* Table S3).

**Table S1.** 11ntR variants with greater specificity for the GUAA TL versus GAAA TL. Specificity values are the median and 95% confidence interval for the difference $\Delta G_{bind,GUAA}$ - $\Delta G_{bind,GAAA}$. Only four mutations, 3C7A, 3C6U, 3G8C, and 3C8G had significant preferences for the GUAA tetraloop.

| Variant | Specificity (kcal/mol) |
|---|---|
| **3C8G** | -0.71 (-0.45, -0.96) |
| **3C7A** | -0.63 (-0.39, -0.89) |
| **3C6U** | -0.64 (-0.11, -1.21) |
| **3G8C** | -0.39 (-0.01, -0.73) |

**Table S2.** The synergistic double mutants. ΔΔΔ*G* values are calculated as described in *SI Appendix* Fig. S6; $\Delta G_{fold}$ and secondary structures are calculated by RNAfold (6).

| Variant | ΔΔΔ*G* (kcal/mol) | $\Delta G_{fold}$ (kcal/mol) | Structure (RNAfold) |
|---|---|---|---|
| **7G6C** | -2.09 (-3.28, -1.38) | -28.5 | (..((_))...) |
| **7G5C** | -1.87 (-2.36, -1.64) | -26.9 | ...((_)).... |
| **5U6A** | -1.49 (-1.76, -1.18) | -30.5 | (((((_)).))) |
| **9A4G** | -1.37 (-2.00, -0.90) | -29.6 | (.(((_)))..) |
| **1G5U** | -1.13 (-1.49, -0.92) | -28.2 | (((((_.))))) |
| **1G4C** | -1.02 (-1.37, -0.82) | -22.7 | (..(._.)...) |
| **5C6A** | -0.99 (-1.44, -0.71) | -26.8 | (..((_))...) |
| **9G4G** | -0.99 (-1.42, -0.77) | -29.6 | (.(((_)))..) |
| **1U5U** | -0.94 (-1.32, -0.70) | -28.2 | (((((_.))))) |
| **5C6C** | -0.83 (-1.23, -0.60) | -25.8 | ...((_)).... |
| **9G11A** | -0.78 (-1.14, -0.57) | -24.6 | (((((_)))..) |
| **1G5G** | -0.73 (-1.19, -0.38) | -24.2 | (.(((_.))).) |
| **9G4U** | -0.71 (-1.17, -0.44) | -30.0 | (((((_)))).) |
| **9A4U** | -0.69 (-0.99, -0.37) | -30.3 | (((((_)))).) |
| **7G2A** | -0.64 (-1.29, -0.23) | -23.7 | ..(((_).)).. |
| **11A4U** | -0.63 (-1.09, -0.34) | -20.0 | (..(._.)...) |
| **7G5U** | -0.55 (-1.05, -0.25) | -28.8 | .((((_)).)). |
| **7G9A** | -0.54 (-1.10, -0.03) | -27.5 | ..(((_)))... |
| **9C4U** | -0.52 (-1.16, -0.11) | -26.8 | (..((_))...) |
| **5G6A** | -0.52 (-0.99, -0.11) | -27.3 | (..((_))...) |
| **7A11A** | -0.51 (-0.95, -0.30) | -19.6 | ...(._.).... |
| **11A4G** | -0.50 (-0.96, -0.19) | -20.0 | (..(._.)...) |
| **7C1U** | -0.47 (-0.70, -0.26) | -25.7 | (..((_))...) |
| **1U5G** | -0.46 (-0.94, -0.05) | -24.7 | (..((_))...) |
| **7G11C** | -0.45 (-0.93, -0.16) | -19.8 | ...(._.).... |
| **7G6A** | -0.39 (-0.86, -0.16) | -26.9 | ...((_)).... |
| **1A5C** | -0.39 (-0.75, -0.18) | -24.2 | (.(((_.))).) |
| **1G5C** | -0.37 (-0.72, -0.16) | -24.2 | (.(((_.))).) |
| **7G11A** | -0.34 (-0.83, -0.12) | -19.8 | ...(._.).... |
| **11A4C** | -0.34 (-0.70, -0.15) | -21.7 | (.(((_..)))) |
| **7G1G** | -0.33 (-0.81,-0.10) | -23.7 | ..(((_.))).. |
| **7A1G** | -0.31 (-0.75, -0.08) | -22.4 | ..(((_.))).. |
| **7G11U** | -0.28 (-0.76, -0.04) | -19.8 | ...(._.).... |
| **11A5U** | -0.27 (-0.64, -0.05) | -23.6 | ((((._..)))) |

**Table S3.** The 11ntR variants whose abundances are significantly over or underpredicted by an exponential relationship with their stabilities. ΔΔ*G* values are reported for two ionic conditions, and "z-score" refers to the residuals' z-scores (Bonferroni-corrected *p*-value of 0.05, see *Methods*). Significantly overpredicted z-scores are in green; underpredicted in red.

| **Variant** | **Abundance** | **ΔΔG (kcal/mol)** 30 mM $Mg^{2+}$ | **z-score** 30 mM $Mg^{2+}$ | **ΔΔG (kcal/mol)** 5 mM $Mg^{2+}$ + 150 mM $K^{+}$ | **z-score** 5 mM $Mg^{2+}$ + 150 mM $K^{+}$ |
|---|---|---|---|---|---|
| **1U9C** | 0 | 1.6 (1.25, 1.75) | -2.65 | 1.47 | -4.06 |
| **1G** | 0 | 0.82 (0.48, 0.97) | -4.47 | 0.65 | -5.37 |
| **1G6C** | 0 | 0.93 (0.53, 1.17) | -4.23 | 2.83 | -1.87 |
| **1G9A** | 0 | 2.12 (1.77, 2.29) | -1.42 | 1.64 | -3.78 |
| **1G9C** | 0 | 1.68 (1.33, 1.84) | -2.45 | 1.37 | -4.22 |
| **1G11C** | 65 | 0.82 (0.62, 0.95) | 3.75 | 0.83 | 2.64 |
| **4G5U** | 0 | 0.32 (-0.11, 0.5) | -5.64 | 1.28 | -4.36 |
| **6U7A** | 0 | 1.1 (0.54, 1.3) | -3.81 | 2.58 | -2.27 |
| **6C** | 42 | 0.32 (-0.07, 0.51) | 1.74 | 2.16 | 3.97 |
| **6C7A** | 0 | 1.1 (0.76, 1.25) | -3.82 | 2.58 | -2.27 |
| **7G** | 1 | 0.31 (-0.18, 0.48) | -4.32 | 1.76 | -2.32 |
| **7G9C** | 0 | 0.3 (-0.06, 0.46) | -5.71 | 2.12 | -3.01 |

**Dataset S1 (separate file).** Binding data for 11ntR variants in this study. "ΔΔG_GAAA" refers to the difference $\Delta G_{\mathrm{bind,mut}}$ - $\Delta G_{\mathrm{bind,wild\ type}}$ binding to the GAAA tetraloop; "ΔΔG_GUAA" refers to the same but binding to the GUAA tetraloop; "ΔΔG_Specificity" refers to the difference $\Delta G_{\mathrm{bind,GUAA}}$ - $\Delta G_{\mathrm{bind,GAAA}}$. Reported $\Delta\Delta G$ values are the median and 95% confidence intervals, calculated as described in the *Methods*; variants which resulted in limits are denoted as such, and their confidence intervals therefore lack an upper bound.

**Dataset S2 (separate file).** Energetic connectivity for 11ntR double mutants binding to a GAAA tetraloop. "Connectivity Type" refers to the four classes defined in Fig. 5. Free energy values are represented by their median and 95% confidence interval and are in units of kcal/mol. "ΔΔΔG" refers to the difference described in *SI Appendix* Fig. S6; "ΔΔG_Double" refers to the $\Delta\Delta G$ value of the double mutant, ΔΔG_Sum_Singles refers to the sum of the two single mutants' $\Delta\Delta G$ values, "ΔΔG_Single_1" refers to the larger of the two single $\Delta\Delta G$ values. p_S, p_C, and p_R refer to adjusted p-values for tests of synergistic, cooperative, and rescue effects, respectively (described in the *Methods*).

**Dataset S3 (separate file).** Thermodynamic fingerprint information for 11ntR variants in this study. Only variants having $\Delta G_{\mathrm{bind}}$ values for three or more scaffolds are listed. "GAAA Variance" refers to the median and 95% confidence interval for the variance in $\Delta\Delta G$ values between the mutant • GAAA and wild-type • GAAA fingerprints, "GAAA p-value" refers to the Benjamini–Hochberg adjusted *p*-value that there is a positive correlation, and "GAAA # non-limit" refers to the number of scaffolds for which $\Delta G_{\mathrm{bind}} < -7.1$ kcal/mol (described in the *Methods*). This information is presented for the correlation between the mutant • G**U**AA and wild-type • G**A**AA fingerprints as well, prefixed by "GUAA" rather than "GAAA."
